# Loss of p32 triggers energy deficiency and impairs goblet cell differentiation in ulcerative colitis

**DOI:** 10.1101/2020.12.09.417915

**Authors:** Annika Sünderhauf, Maren Hicken, Heidi Schlichting, Kerstin Skibbe, Mohab Ragab, Annika Raschdorf, Misa Hirose, Holger Schäffler, Arne Bokemeyer, Dominik Bettenworth, Anne G. Savitt, Sven Perner, Saleh Ibrahim, Ellinor I. Peerschke, Berhane Ghebrehiwet, Stefanie Derer, Christian Sina

## Abstract

Cell differentiation in the colonic crypt is driven by a metabolic switch from glycolysis to mitochondrial oxidation. Mitochondrial and goblet cell (GC) dysfunction have been attributed to the pathology of ulcerative colitis (UC). We hypothesized that p32/gC1qR/HABP1, which critically maintains oxidative phosphorylation, is involved in GC differentiation and hence in the pathogenesis of UC. In UC patients in remission, colonic GC differentiation was significantly decreased compared to controls in a p32-dependent manner. Plasma/serum lactate and colonic pAMPK level were increased, pointing at high glycolytic activity and energy deficiency. Consistently, *p32* silencing in mucus-secreting HT29-MTX cells abolished butyrate-induced differentiation and induced a shift towards glycolysis. In mitochondrial respiratory chain complex V-deficient mice, colonic p32 expression correlated with loss of differentiated GCs, resulting in a thinner mucus layer. Conversely, feeding mice an isocaloric glucose-free, high-protein diet increased mucosal energy supply that promoted colonic p32 level, GC differentiation and mucus production. We here describe a new molecular mechanism linking mucosal energy deficiency in UC to impaired p32-dependent GC differentiation that may be therapeutically prevented by nutritional intervention.

## INTRODUCTION

Ulcerative colitis (UC), as one main phenotype of inflammatory bowel disease (IBD), is a chronic, relapsing-remitting immune mediated disorder of the human gastrointestinal tract, in which inflammation is localized to the large intestine and restricted to the mucosa. While the exact pathophysiology is still not fully understood, genetic, environmental and immune-mediated factors contribute to disease onset and recurrence in UC. Loss of intestinal epithelial and mucus barrier integrity leading to bacterial translocation is commonly accepted as a major cause of inflammation (1). Correct cellular differentiation, which is pivotal for the cryptic architecture and thus barrier integrity, is a highly energy demanding process (2), strongly suggesting that mitochondrial dysfunction plays a key role in both the onset and recurrence of the disease. Of main interest, mitochondrial dysfunction in epithelial cells, defective goblet cell differentiation and mucus depletion in UC have been independently reported in several studies (3-9). Nevertheless, mechanistic evidence linking cellular energy metabolism to goblet cell differentiation and UC pathogenesis is still missing.

Cells of the colonic mucosa utilize different mechanisms to maintain their energy homeostasis. Energy generation in cells of the lower third of the crypt (*e*.*g*. intestinal stem cells) mainly depend on gly-colysis, while short chain fatty acids (SCFA) inhibit stem/progenitor cell proliferation (10, 11). In contrast, differentiated post-mitotic cells of the upper third of the crypt (*e*.*g*. goblet cells) maintain their energy level through mitochondrial *β*-oxidation of SCFA such as butyrate as well as long-chain fatty acids and the oxidative phosphorylation (OXPHOS) system (2, 11-13). We recently reported a cellular mechanism, whereby caspase-1 dependent cleavage of p32, a protein that critically maintains OXPHOS function, induces a metabolic shift from mitochondrial OXPHOS to cytosolic aerobic glycolysis (14). This metabolic shift led to an enhancement of cell proliferation and a decrease in cell differentiation of cancer cells and is potentially involved in the transition of transient amplifying cells into post mitotic cells.

In the intestinal crypt, differentiation of goblet cells occurs along the metabolic trajectory of shifting energy source. Secretory precursor cells in the transit amplifying zone are characterized by high expression of atonal basic helix-loop-helix transcription factor 1 (*ATOH1*, also referred to as *HATH1* in humans and *Math1* in mice) (15) and further by high level of SAM pointed domain-containing Ets transcription factor (*SPDEF1*) (16). Kruppel-like factor 4 (*KLF4*) expressing, terminally differentiated goblet cells are particularly specialized in the production and secretion of highly glycosylated proteins, so called mucins, with mucin 2 (MUC2) being the most abundant in the colon and small intestine. Notably, *klf4*-deficient mice display defective goblet cell differentiation with a decrease of about 90% of colonic goblet cells (17). Reduced numbers of goblet cells in line with colonic mucus depletion have been suggested as histological hallmarks of UC (6). Gersemann *et al*. showed, that induction of goblet cell differentiation during inflammation is impaired in UC but not in Crohn’s disease (CD) (8). Furthermore, differentiation defects of intestinal stem cells have been found to be accompanied by a barrier dysfunction, leading to intestinal inflammation and/or cancer development (18).

The OXPHOS system has been found to be highly active in goblet cells (2). Therefore, differentiated goblet cells are expected to be highly affected by reduced OXPHOS activity. Supporting this hypothesis, we recently published that loss of OXPHOS-stabilizing p32 by inflammasom-driven cleavage reduces goblet cell differentiation state, *in vitro* (14).

In 1980, Roediger *et al*. hypothesized, that pathogenesis of UC is linked to energy-deficiency. More specifically, the authors found reduced butyrate oxidation rates in isolated colonocytes from UC patients compared to healthy controls (3). Furthermore, two independent studies reported reduced mitochondrial respiratory chain complex activity accompanied by mucosal ATP depletion in UC patients (4, 5). Interestingly, alterations in all three studies were already present in non-inflamed tissue, implicating mitochondrial dysfunction as a pathophysiological cause rather than consequence in UC.

The ubiquitous nuclear encoded multi-functional protein p32 critically maintains OXPHOS and was independently identified as a subunit of the human pre-mRNA splicing factor SF2 (19, 20), as complement component 1q binding protein (C1qbp, gC1qR) (21) and as hyaluronic acid binding protein (HABP1) (22). P32 expression is integral to mitochondrial energy maintenance, with energy generation *via* OXPHOS being nearly absent in *p32* knockout cells (14, 23) and *p32*-deficient mice being embryonic lethal (24). Cumulative data indicate that one of the major functions of p32 is to maintain mitochondrial function by regulating mitochondrial protein translation (24, 25).

Taken together, we hypothesized p32 to be involved in the maintenance of the metabolic trajectory within the intestinal crypt, thereby enabling the metabolic switch from glycolysis to mitochondrial OXPHOS, which is necessary for terminal differentiation of intestinal stem cells towards goblet cells. Since mitochondrial dysfunction and defects in goblet cell differentiation have been attributed to UC pathogenesis, we aimed at investigating colonic expression of p32 in UC patients, as well as studying mechanistic backgrounds and possible modulation of OXPHOS driven goblet cell differentiation.

## RESULTS

### UC patients in remission display decreased colonic p32 expression, increased glycolysis and cellular energy deficiency

Stem cells in the lower part of the colonic crypt are mainly dependent on glycolysis, while there is a gradient towards an increase in energy generation *via* OXPHOS and a decrease in glycolysis towards the differentiated cells at the tip of the crypt (2, 10, 12). Cellular differentiation occurs alongside this gradient of shifting energy source (**Figure 1A**) and we postulated p32 as a main driver of mitochondrial OXPHOS to be involved in its maintenance. Indeed, p32 is highly expressed in the upper part of the colonic crypt, together with mitochondrial marker TOMM22 and goblet cell differentiation marker KLF4 (**Figure 1B** and **C**). When *p32* mRNA expression was analyzed in a cohort of intestinal biopsies (ileum to sigmoidal colon) from 15 non-IBD controls and 22 UC patients in remission, we found reduced *p32* level in UC patients (**Table 1** and **Figure 1D**). Expression data from ileal and colonic biopsies were combined, since *p32* mRNA expression did not differ between ileal and colonic tissue within patients (**Figure 1E**). At least two isoforms of *p32* have been described, one encoding and one lacking the mitochondrial leader sequence in exon 1 (26). Hence, we investigated exon expression in a subset of intestinal biopsies from 10 non-IBD controls and 9 UC patients in remission. Expression of all exons, and therefore most likely all isoforms, of the *p32* transcript were reduced in the intestine of UC patients compared to non-IBD controls (**Figure 1F** and **Supplementary Figure 1A**).

**Figure 1:**
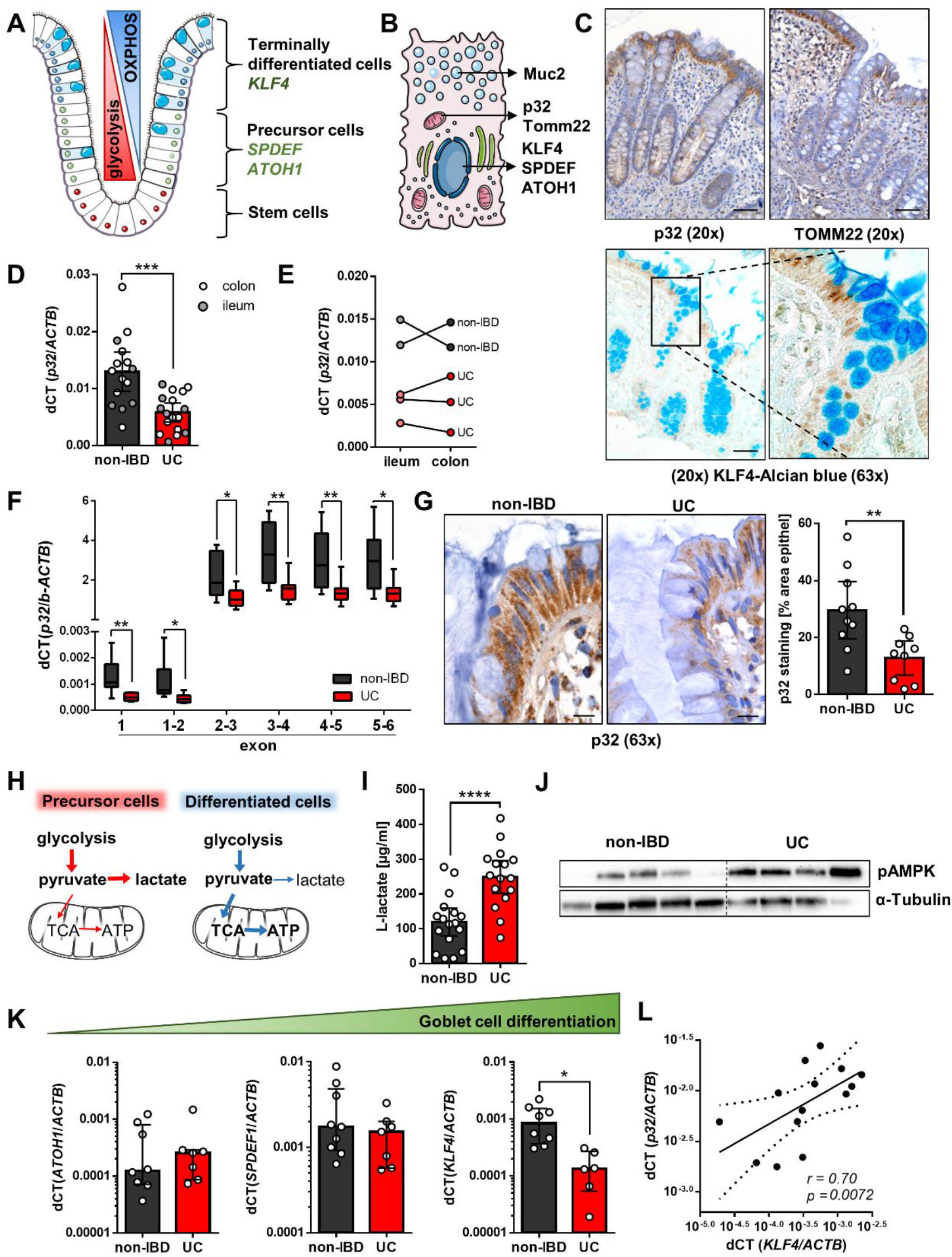
UC patients in remission display reduced p32 level, mucosal energy deficiency and impaired goblet cell differentiation. **A)** A model for energy generation and goblet cell differentiation in the colonic crypt and **B)** schematic subcellular localization of proteins of interest were generated by modifying images from Servier Medical Art (52). **C)** Representative IHC staining of p32 (clone EPR8871), Tomm22 and KLF4 in human colonic biopsies. Scale bar = 50 µM. Expression of transcripts of interest was measured by qRT-PCR in **D)** and **E)** ileal and colonic biopsies and **K)** colonic biopsies from non-IBD and UC patients in remission. **F)** *P32* exon expression was analyzed by taqman assay. Non-IBD: n = 10; UC: n = 7 and 6 for exon 1 and exon 1-2, respectively and n = 8-9 for all other exon junctions. **G)** Representative IHC staining and corresponding quantification of p32 (clone EPR8871) expression in the upper part of the colonic crypt in biopsies from non-IBD controls and UC patients in remission. Scale bar = 10 µM **H)** Schematic visualization of energy generation in the intestinal crypt. **I)** L-lactate level were measured in serum or plasma samples and **J)** WB experiments were performed in colonic biopsies from non-IBD controls and UC patients in remission. **L)** Colonic *p32* mRNA expression was correlated against *KLF4* mRNA expression in non-IBD controls and UC patients in remission. **D), F)** and **K)** Unpaired t-test with Welch’s correction; **G)** and **I)** Unpaired t-test; **L)** Spearman’s rank correlation coefficient; results are shown as **D), G)** and **I)** mean ± 95% CI; **F)** Box & whiskers plot min to max; **K)** median ± interquartile range; * p ≤ 0.05, ** p ≤ 0.01, *** p ≤ 0.001, **** p ≤ 0.0001.

**Table 1:** Patients’ characteristics native and paraffin-embedded biopsies.

|  | P32 mRNA expression |  | IHC analysis |  |  |
| --- | --- | --- | --- | --- | --- |
| Total number of patients included | 38 |  | 24 |  |  |
| Non-IBD | 15 (39.5%) |  | 10 (41.7%) |  |  |
| UC remission non-inflamed | 23 (60.5%) |  | 9 (37.5%) |  |  |
| UC inflamed | - |  | 5 (20.8%) |  |  |
| Gender ratio women : men (unknown) |  |  |  |  |  |
| Non-IBD | 7 (46.7%) : 7 (46.7%) (1 (6.7%)) |  | 5 (50.0%) : 5 (50.0%) |  |  |
| UC remission non-inflamed | 12 (52.2%) : 11 (47.8%) |  | 7 (77.8%) : 2 (22.2%) |  |  |
| UC inflamed | - |  | 3 (60.0%): 2 (40.0%) |  |  |
| Mean age ± SD (unknown) |  |  |  |  |  |
| Non-IBD | 61.4 ± 17.9 (1) |  | 53.4 ± 11.3 |  |  |
| UC remission non-inflamed | 50.2 ± 16.6 (1) |  | 60.0 ± 13.5 |  |  |
| UC inflamed | - |  | 51.6 ± 7.7 |  |  |
| Origin of biopsies | Non-IBD | UC | Non-IBD | UC non-inflamed | UC inflamed |
| Ileum | 4 (26.7%) | 5 (21.7%) | - | - | - |
| Caecum | - | 4 (17.4%) | - | - | - |
| C. ascendence | 2 (13.3%) | 2 (8.7%) | - | - | - |
| Flexura hepatica | 1 (6.7%) | - | - | - | - |
| C. transversum | 1 (6.7%) | 1 (4.3%) | 1 (10.0%) | - | - |
| C. descendence | - | 1 (4.3%) | 3 (30.0%) | 3 (33.3%) | 1 (20.0%) |
| C. sigmoideum | 6 (40.0%) | 9 (39.1%) | 6 (60.0%) | 6 (66.7%) | 4 (80.0%) |
| Rectum | 1 (6.7 %) | 1 (4.3%) | - | - | - |
| Colon unclassified | - | - | 1 (10.0%) | 1 (11.1%) | - |
| Medication | Non-IBD | UC | Non-IBD | UC non-inflamed | UC inflamed |
| Mesalazine/Mesalazine klysmen | - | 12 (52.2%) | - | 5 (55.6%) | 2 (40.0%) |
| Prednisone | - | 6 (26.1%) | 1 (10.0%) | - | 2 (40.0%) |
| Azathioprine | - | 5 (21.7%) | - | - | - |
| Sulfasalazine | - | 2 (8.7%) | - | 1 (11.1%) | - |
| Tacrolimus | - | 2 (8.7%) | - | - | 1 (20.0%) |
| Budesonide klysmen | - | - | - | - | 2 (40.0%) |
| Metronidazole | - | - | 1 (10%) | - | 1 (20.0%) |
| Sirolimus | - | 1 (4.3%) | - | - | - |
| Hydrocortisone rectal foam | - | 1 (4.3%) | - | - | - |
| Olsalazine | - | 1 (4.3%) | - | - | - |
| Ciprofloxacin | 1 (6.7%) | - | 3 (30.0 %) | 1 (11.1 %) | 1 (20.0 %) |

Due to the fact that mitochondrial function is highly affected by ageing (27) and various therapeutic regiments, we related *p32* mRNA expression to patients’ age and tested for potential influences of commonly prescribed therapeutics within our cohort such as prednisolone, mesalazine and azathioprine. *P32* mRNA expression did not correlate with age in either non-IBD controls or UC patients (**Supplementary Figure 1B**). In line with previous studies, which showed azathioprine to impair cell proliferation (28), azathioprine treatment was associated with higher *p32* mRNA level in UC patients, an effect neither observed under mesalazine nor prednisolone therapy (**Supplementary Figure 1C** and **D**). Therefore, biopsies from patients receiving azathioprine treatment were excluded from data presented in Figure 1 and 2.

**Figure 2:**
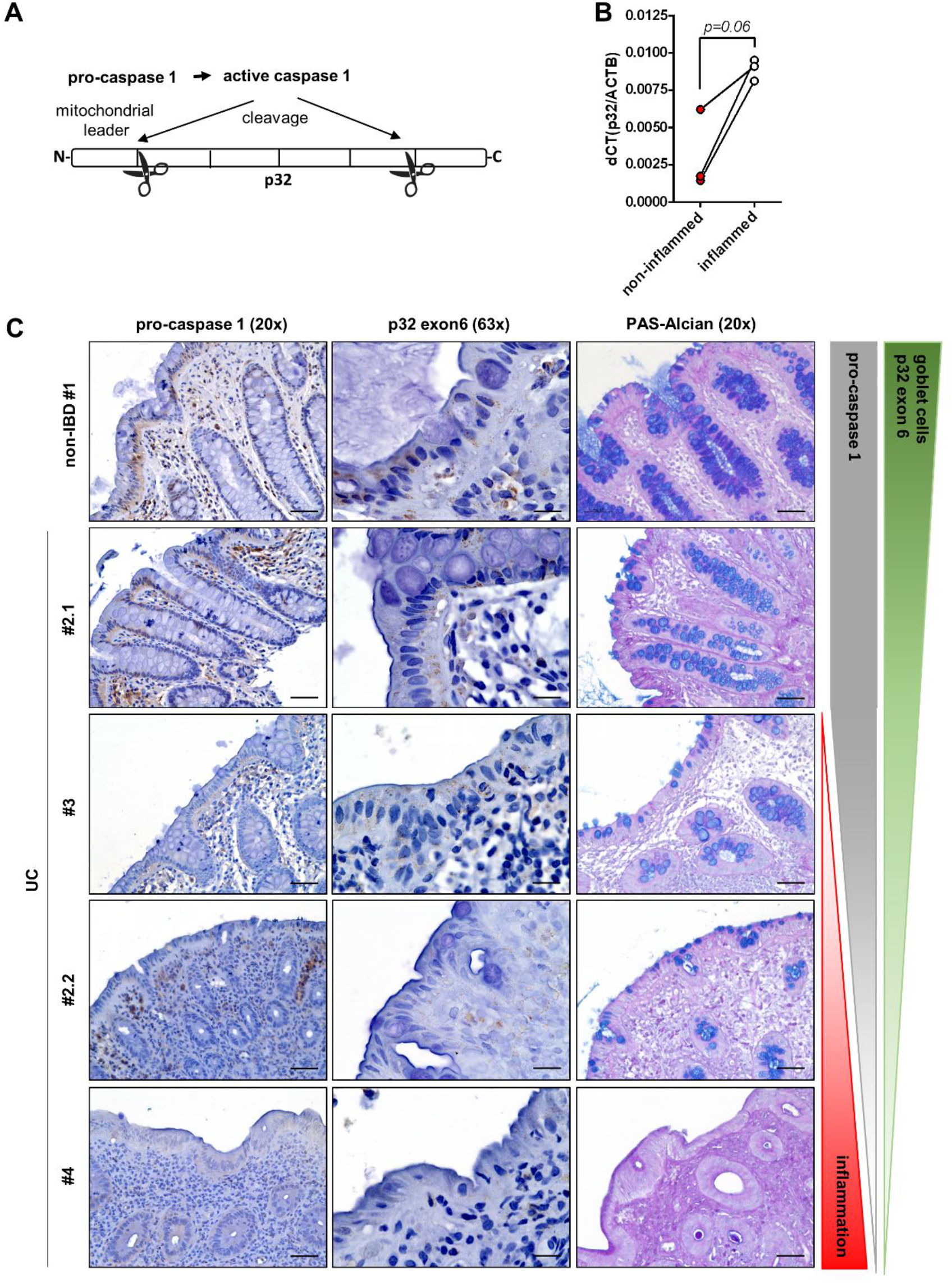
Goblet cell loss correlates with inflammasome activation and decrease of full-length p32 level in active UC. **A)** Schematic visualization of p32 cleavage by active Caspase-1. **B)** P32 mRNA expression in paired biopsies from non-inflamed and inflamed intestinal tissue sections were quantified by qRT-PCR. **C)** Representative IHC staining against pro-caspase1 and p32 exon 6 as well as PAS-Alcian staining in tissue biopsies from the descending colon or sigma; #1: non-IBD non-inflamed; #2.1: UC non-inflamed; #3 UC low grade inflammation; #2.2 UC medium-grade inflammation; #4 UC high grade inflammation. Representative images from 8 biopsies each categorized as non-IBD controls or UC non-inflamed and 5 UC inflamed samples are displayed. Scale bar = 50 µM **B)** Paired t-test.

To investigate whether differences in *p32* mRNA level are also reflected on protein level, a set of ten colonic biopsies collected from non-IBD patients was compared to nine colonic biopsies collected from UC patients in remission *via* IHC staining of the p32 protein (clone EPR8871). P32 staining was densitometrically quantified in the upper third of the crypt and revealed a significantly lower p32 positive area in UC patients compared to non-IBD controls (**Table 1** and **Figure 1G**). Further, UC patients displayed increased L-lactate level in plasma/serum samples compared to non-IBD controls as well as high phosphorylation of AMPK in colonic biopsies, pointing to increased glycolysis activity and mucosal energy deficiency in UC patients. (**Table 2** and **Figure 1H-J**).

**Table 2:** Patients’ characteristics serum samples and western blot biopsies.

|  | Serum/plasma | WB analysis |  |
| --- | --- | --- | --- |
| **Total number of patients included** | **33** | **9** |  |
| Non-IBD | 17 (51.5%) | 5 (55.6%) |  |
| UC remission | 16 (48.5%) | 4 (44.4%) |  |
| **Gender ratio women : men** |  |  |  |
| Non-IBD | 9 (52.9%) : 8 (47.1%) | 3 (60.0%) : 2 (40%) |  |
| UC remission | 7 (43.8%) : 9 (56.3%) | 1 (25%) : 3 (75%) |  |
| **Mean age ± SD (unknown)** |  |  |  |
| Non-IBD | 30.3 ± 9.7 | 49.8 ± 24.2 |  |
| UC remission | 43.8 ± 7.7 (10) | 60.15 ± 17.04 |  |
| **Origin of biopsies** | Not applicable | **Non-IBD** | **UC** |
| Caecum |  | - | 1 (25.0%) |
| C. ascendence |  | 1 (20.0%) | 1 (25.0%) |
| Flexura hepatica |  | 1 (20.0%) | -- |
| C. descendence |  | 1 (20.0%) | 1 (25.0%) |
| C. sigmoideum |  | 1 (20.0%) | - |
| Rectum |  | 1 (20.0%) | 1 (25.0%) |
| **Medication** | **UC** | **UC** |  |
| Mesalazine/Mesalazine klysmen | 15 (93.8%) | 2 (50%) |  |
| TNF-α inhibitors | 8 (50.0%) | - |  |
| Prednisone | 1 (6.3%) | 2 (50%) |  |
| Vedolizumab | 3 (18.8%) | - |  |
| Budesonide klysmen | 2 (12.5%) | - |  |
| Thiopurines | 2 (12.5%) | - |  |
| Hydrocortisone rectal foam | - | 1 (25%) |  |
| Ciprofloxacin | - | 1 (25%) |  |
| Sulfasalazine | - | 1 (25%) |  |
| Tacrolimus | - | 1 (25%) |  |

### Colonic goblet cell differentiation is impaired in UC patients in remission and goblet cell number decreases with increasing degree of inflammation

High glycolytic activity characterizes cell metabolism in proliferating precursor cells rather than in non-dividing differentiated cells (10). Since goblet cell function has been previously proposed to be impaired in UC (3, 7, 8), we focused on analyzing differentiation status of this cell entity. Interestingly, expression of terminal goblet cell differentiation marker *KLF4* was significantly downregulated in colonic biopsies (hepatic flexure to sigmoid colon) from UC patients in remission compared to non-IBD controls (**Figure 1K**). Additionally, colonic *KLF4* mRNA expression significantly correlated with *p32* mRNA expression (**Figure 1L**), supporting the hypothesis, that impaired terminal goblet cell differentiation in UC is a result of defective energy generation *via* p32-driven OXPHOS. Meanwhile, transcript levels of goblet cell precursor markers *ATOH1* and *SPDEF1* were not statistically different (**Figure 1K**).

In the next set of experiments, we analyzed p32 expression and goblet cell appearance in non-inflamed and inflamed tissue sections of UC patients in remission or active disease. Inflammasomes, as part of the innate immune system, are responsible for the initiation of inflammatory responses, mediated by the activation of caspase-1 among others (29). We have recently published, that active caspase-1 cleaves p32 at two distinct sites (exon 1-2 junction and in exon 5), thereby preventing mitochondrial import of p32. This mechanism results in a shift in energy generation of tumor cells from OXPHOS towards aerobic glycolysis (**Figure 2A**) and abrogation of differentiation of goblet cell-like HT29-MTX cells, *in vitro* (14). On mRNA level, *p32* was upregulated in inflamed tissue biopsies compared to samples from non-inflamed regions within patients with active UC (**Figure 2B**). Of note, protein expression of pro-caspase 1 was reduced in inflamed but not non-inflamed tissue areas of UC patients indicating inflammasome activation in respective regions. Consistent with reported caspase-1 induced p32 cleavage, binding of an antibody against p32 exon 6 was reduced in UC inflamed tissue sections compared to non-IBD controls in a disease activity dependent manner. Furthermore, blinded evaluation of PAS-Alcian blue staining revealed reduced staining intensity of goblet cell granules in UC non-inflamed tissue compared to non-IBD controls under basal conditions. The amount of mucus-filled goblet cells was reduced under low grade inflammation and further decreased with increasing degree of mucosal inflammation (**Table 1, Figure 2D** and **Supplementary Figure 2**). Overall, these findings support our previous observation that caspase-1 cleavage of p32 leads to abrogation of goblet cell differentiation (14), thereby further reducing mitochondria-localized and functional p32 and differentiated goblet cells in UC.

### Goblet cell differentiation is dependent on OXPHOS and p32 *in vitro*

To test our hypothesis that OXPHOS-driven goblet cell differentiation in the intestinal crypt is dependent on p32, we next screened a range of human colorectal carcinoma cell lines for expression of goblet cell differentiation markers, *MUC2, Mucin5AC* (*MUC5AC)* and *p32*. HT29-MTX cells depicted high basal mRNA level of *SPDEF1*, indicating a goblet cell precursor phenotype as well as *MUC5AC* but not *MUC2*. While DiFi cells showed high levels of both *ATOH1* and *KLF4*, the analyses of T84 cells indicated terminal differentiation reflected by high expression of *KLF4* and *MUC2*. All these three goblet cell-like cell lines similarly expressed *p32* mRNA (**Supplementary figure 3A**). To find an inducible cell line model to study dependency of goblet cell differentiation on mitochondrial activity in vitro, β-oxidation and hence OXPHOS in HT29-MTX, T84 and DiFi cells was boosted through stimulation with the short-chain fatty acid butyrate in the presence or absence of the proinflammatory stimulus LPS, frequently present in the intestine (**Figure 3A** and **B**). Butyrate stimulation induced terminal goblet cell differentiation of HT29-MTX but not of T84 or DiFi cells, reflected by induction of KLF4 expression, which was abrogated in the presence of LPS (**Figure 3B** and **G**). Butyrate-triggered terminal goblet cell differentiation of HT29-MTX cells was accompanied by an increase in oxygen consumption rate (OCR) but not in extracellular acidification rate (ECAR) (**Figure 3C**) (14), underlining the importance of a metabolic switch towards OXPHOS in goblet cell differentiation. Furthermore, differentiated HT29-MTX cells displayed increased mucin granule formation, decreased cell proliferation and enhancement of secreted Muc5AC (**Figure 3D-G)**. Of note, *p32* mRNA expression was not altered upon butyrate stimulation (**Supplementary Figure 3B**). To test whether goblet cell differentiation is indeed dependent on p32, we performed siRNA-induced silencing experiments in HT29-MTX cells. Of main interest, induction of goblet cell differentiation *via* butyrate was abrogated in *p32*-silenced HT29-MTX cells accompanied by increased lactate level, indicating a switch in energy metabolism towards aerobic glycolysis. Thus, supporting the idea that p32 maintains mitochondrial function and thereby ensures goblet cell differentiation (**Figure 3H** and **I**). OXPHOS is a lot more efficient in the production of ATP compared to aerobic glycolysis. Therefore, we proposed a pivotal role for cellular energy supplied by the mitochondrial OXPHOS system not only for goblet cell differentiation, but also for mucus secretion. To test this hypothesis, HT29-MTX cells were first terminally differentiated by post-confluent growth (30) **(Supplementary Figure 3C)**, followed by stimulation with OXPHOS complex V blocker oligomycin or the uncoupling agent DNP (**Figure 3J**). As expected, blocking of OXPHOS function by oligomycin resulted in a shift of cellular energy metabolism from OXPHOS to glycolysis **(Figure 3K)**. Moreover, mucus secretion was impaired by oligomycin as well as by DNP, reflected by a dose-dependent downregulation of secreted but not intracellular Muc5AC (**Figure 3L** and **Supplementary Figure 3D** and **E**), supporting the idea that mucus secretion is a highly energy demanding process enabled by efficient OXPHOS activity.

**Figure 3:**
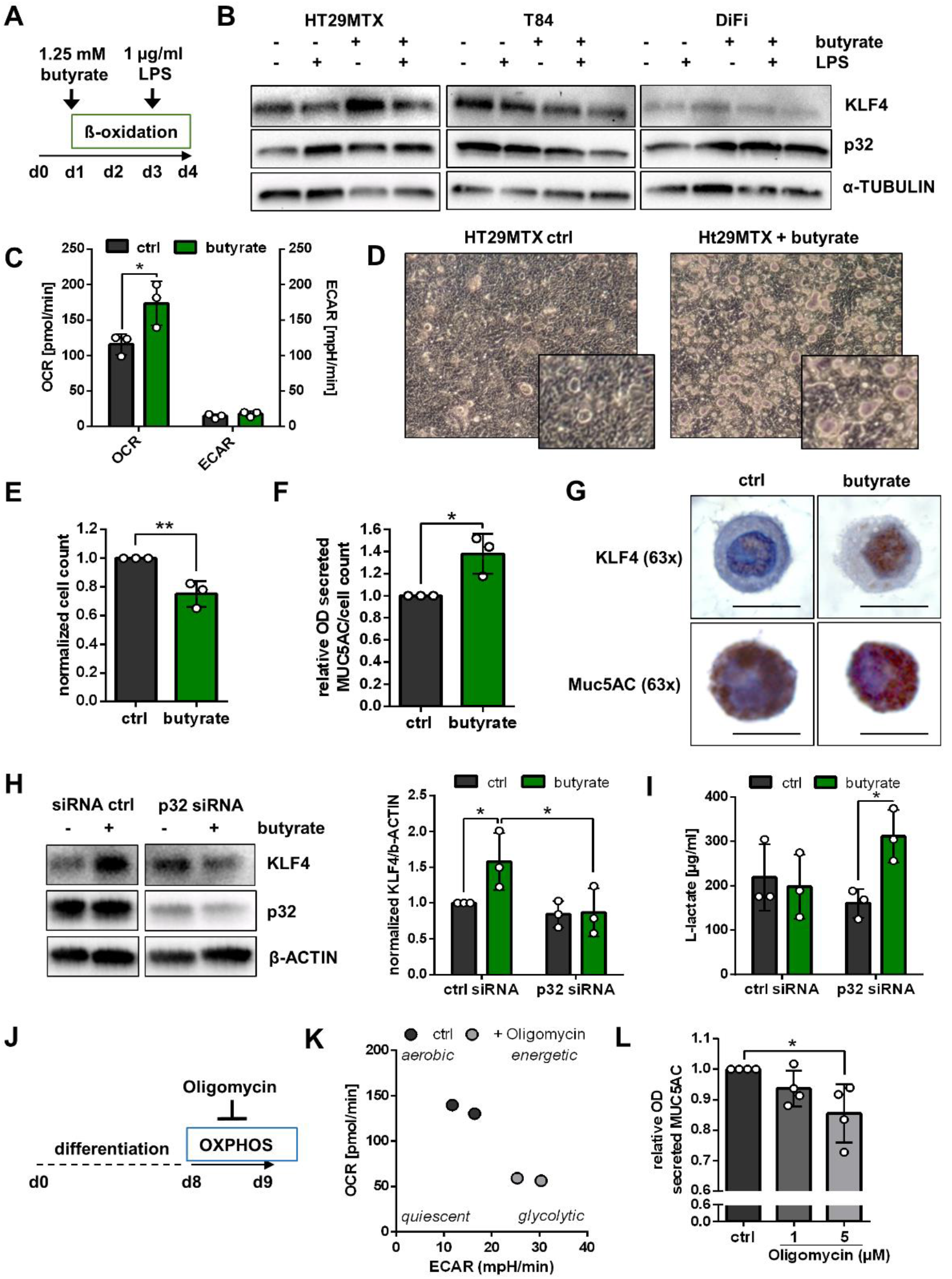
Mucin secretion and goblet cell differentiation is dependent on energy supplied by mitochondrial respiration. **A)** and **J)** Graphical setup of cell culture experiments. **B)** WB experiments were performed from whole protein extracts with respective antibodies in cells stimulated with 1.25 mM butyrate and/or 1 µg/ml LPS (p32 clone EPR8871). **C)** Basal oxygen consumption rate (OCR) and extracellular acidification rate (ECAR) were measured by Seahorse assay (14). **D)** Representative image of HT29-MTX cell growth characteristics. **E)** Cell counts are presented as fold change for each individual experiment. **F)** Muc5AC level in the supernatant were measured by ELISA, normalized to cell count and are displayed as fold change for each individual experiment. **G)** KLF4 and MUC5AC IHC staining was performed in paraffin-embedded butyrate-stimulated or control HT29-MTX cells. Scale bar = 10 µM. **H)** to **I)** For siRNA knockdown, HT29-MTX cells were stimulated with 50 nM p32 siRNA or respective control for 96 hours and butyrate for 72 hours. **H)** Representative WB and quantification from whole protein extracts (p32 clone 60.11) and **I)** L-Lactate level from corresponding cell culture supernatants. **K)** Seahorse measurement of HT29-MTX cells before and after 2 µM oligomycin injection **L)** Muc5AC level in the cell culture supernatant after 24h stimulation with oligomycin was measured by ELISA and normalized to each control. **C)** paired t-test; **E)** and **F)** unpaired t-test; **H)** and **I)** uncorrected Fisher’s test **L)** One-way ANOVA with Bonferroni’s post-hoc test; results are shown from 3 independent experiments with the exception of **K)** n = 2; Data are shown as mean ± SD; * p ≤ 0.05, ** p ≤ 0.01.

### ATP8-mutant mice display low colonic p32 expression in concert with loss of OXPHOS and goblet cells

To investigate the observed UC phenotype of low colonic p32 level, energy deficiency and defective goblet cell differentiation in a mouse model, we applied conplastic respiratoty chain complex V-mutant mice. These mice carry a mutation in the mitochondrial encoded ATP8-synthase resulting in diminished respiratory capacity and ATP production with parallel induction of energy generation *via* non-mitochondrial glycolysis in various cell entities (31-33) (**Figure 4A**). Specifically, we found that ATP8-mutant mice displayed reduced *p32* mRNA expression and diminished p32 protein level especially in differentiated intestinal epithelial cells in the upper part of colonic crypts (**Figure 4B** and **C**), while serum L-lactate levels were similar between strains (**Supplementary Figure 4A**). In line with the phenotype observed in UC patients, loss of p32 in ATP8-mutant mice was associated with altered colonic goblet cell differentiation represented by decreased *klf4* mRNA expression, diminished mucus filling of goblet cells and a reduced thickness of the colonic mucus layer. Further, *KLF4* mRNA expression significantly correlated with *p32* mRNA expression in colonic samples from B6-wt and ATP8-mutant mice (**Figure 4D**-**F**). Expression of *atoh1* and *spdef1* was not altered which was comparable to observations in UC patients (**Figure 4D**). Furthermore, expression of the proliferation marker *ki67* and the stem cell marker *lgr5* were not different in colonic biopsies from ATP8-mutant mice (**Supplementary Figure 4B)**. Taken together, we here present a mouse model with low intestinal p32 level and diminished energy generation via OXPHOS to depict reduced numbers of terminally differentiated goblet cells in the colon, strengthening the notion that especially goblet cell differentiation is highly sensitive to mitochondrial dysfunction.

**Figure 4:**
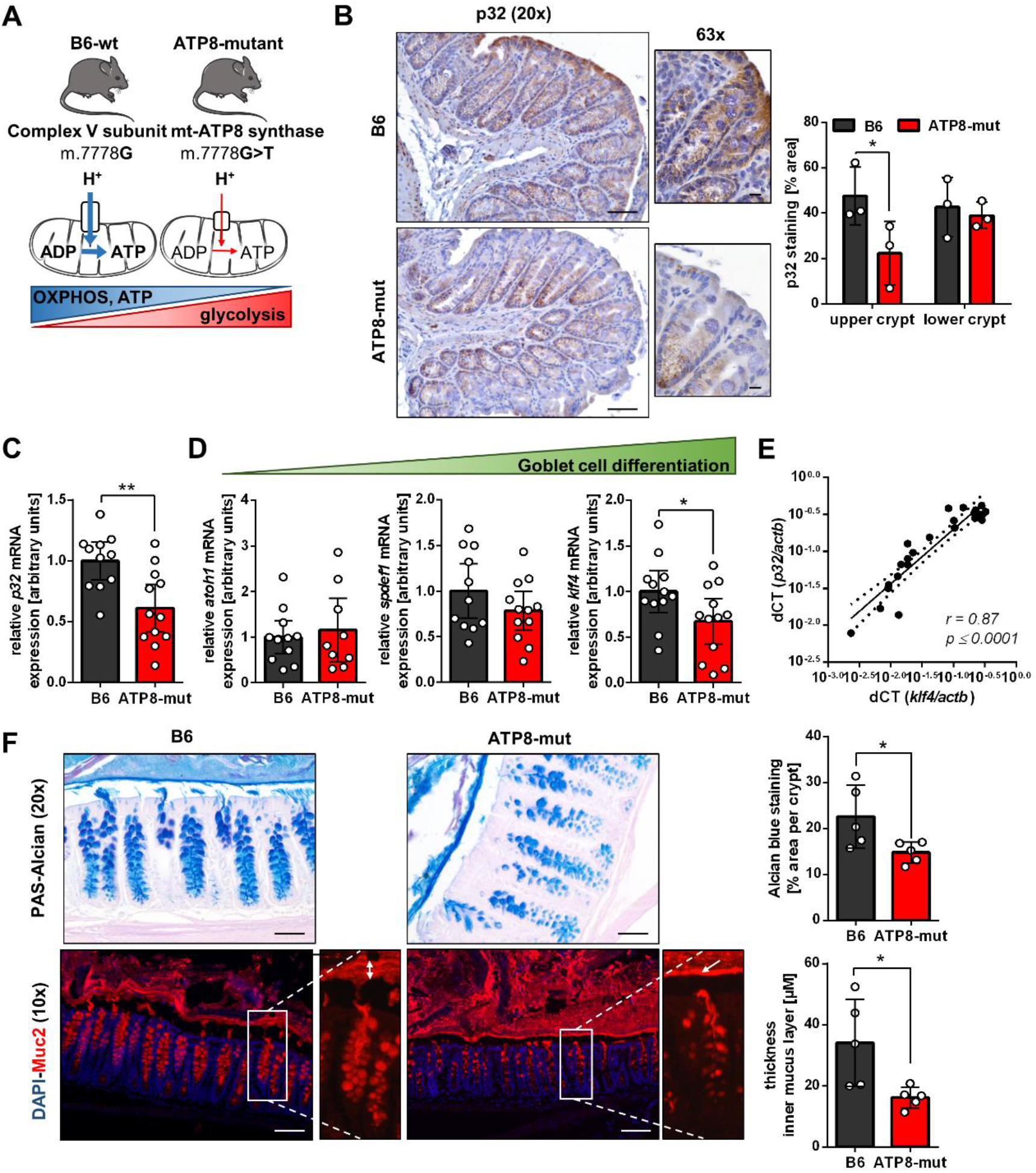
Mitochondrial dysfunction in mice is accompanied by defective goblet cell differentiation. **A)** Schematic overview of the mutation in subunit 8 of the ATP-synthase in ATP8-mut mice and published metabolic imbalance (32, 33). **B)** Representative IHC staining and according quantification of p32 (clone EPR8871) expression in colonic biopsies of B6-wt and ATP8-mut mice (n = 3 mice per group). Scale bar 20x = 50 µM; scale bar 63x = 10 µM. **C)** and **D)** Expression of transcripts of interest was performed by qRT-PCR in colonic biopsies from B6-wt and ATP8-mut mice. Data were normalized to *ß-actin* and are displayed as relative values to B6-wt mice for each sampling round. **E)** Colonic *p32* mRNA expression was correlated against *klf4* mRNA expression in B6-wt and ATP8-mut mice. **F)** Representative PAS-Acian and Muc2 fluorescent staining with according quantification in Carnoy’s fixed colonic tissue samples from B6-wt and ATP8-mut mice. Scale bar PAS-Alcian = 50 µM; scale bar Muc2 IF = 100 µM. Arrow indicates inner mucus layer. **B)** to **D)** Unpaired t-test; **E)** Spearman’s rank correlation coefficient; **F)** unpaired t-test with Welch’s correction; **B)** and **F)** results are shown as mean ± SD; **C)** and **D)** results are shown as mean ± 95% CI; * p ≤ 0.05, ** p ≤ 0.01.

### A glucose-free nutritional intervention promotes colonic p32 expression and goblet cell differentiation in mice

Finally, we aimed to study regulation of goblet cell differentiation *via* the enhancement of intestinal p32 expression in a glucose-free, high-protein nutritional intervention in mice. Since availability of nutrients critically affects cellular metabolism (34), we hypothesized that withdrawal of glucose from the diet and isocaloric replacement of glucose by the protein casein may result in a metabolic shift towards mitochondrial oxidation (**Figure 5A**). Adult C57BL/6 mice were fed a glucose-free high-protein (GFHP) or an isocaloric control diet (ctrl) for an average of 70 days before organ sampling and molecular analysis. Food consumption, body weight, serum glucose and lactate level were similar between diets (**Figure 5B** and **C, Supplementary Figure 4C** and **D**). Of main interest, GFHP diet fed mice exhibited increased p32 protein level in the upper part of the colonic crypt which was not due to elevated *p32* mRNA level (**Figure 5D-F**). Simultaneously, GFHP diet fed mice displayed high colonic energy level reflected by low phosphorylation of AMPK (**Figure 5E**). Eventually, we tested whether enhanced p32 expression would also result in enhanced goblet cell differentiation. In comparison to control mice, increased KLF4 mRNA and protein expression as well as a thicker colonic mucus layer were potent indicators for induction of terminal differentiation of goblet cells under GFHP diet (**Figure 5G and H**). Further, *spdef1* as a marker for secretory progenitor cells was reduced in GFHP diet mice supporting the idea, that p32 expression is pivotal for the transition from secretory precursors towards terminal differentiated goblet cells (**Figure 5G**). Expression of intestinal stem cell marker *lgr5* and proliferation marker *ki67* were unaltered upon GFHP diet (**Supplementary Figure 4E**). Taken together, nutritional intervention by glucose restriction in the presence of high protein intake appears as a promising tool to enhance colonic p32, thereby improving cellular energy supply and finally promoting goblet cell differentiation.

**Figure 5:**
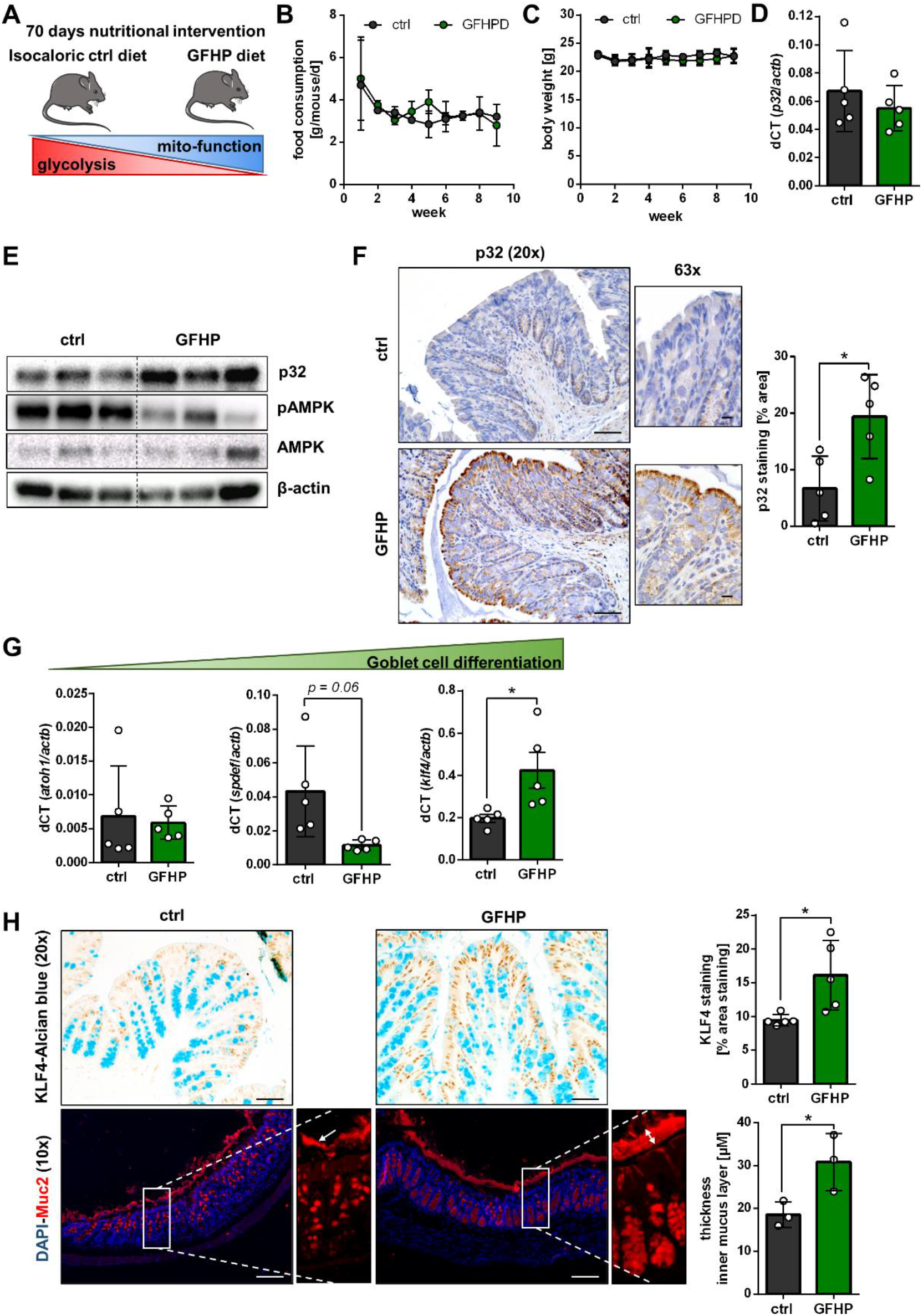
GFHP diet increased mucosal energy supply, induces colonic p32 protein expression and promoted goblet cell differentiation. **A)** Hypothesized metabolic switch upon glucose free high protein (GFHP) dietary intervention in mice. Weekly, **B)** food consumption and **C)** mice body weight was determined (n = 6 from two independent experiments). Expression of transcripts was measured **D)** via taqman probes for p32 exon 3-4 or **G)** by SYBR qRT-PCR in colonic biopsies from ctrl and GFHP diet mice. **E)** WB experiment of whole protein extracts from colonic samples from ctrl and GFHP mice (p32 clone EPR8871). **F)** and **H)** Representative p32 (clone EPR8871) and KLF4-Alcian blue IHC staining of PFA-fixed colonic tissue samples as well as MUC2 fluorescent staining of Carnoy’s fixed tissue are presented with corresponding quantifications. Scale bar 10x = 100 µM; scale bar 20x = 50 µM; scale bar 63x = 10 µM. Arrow indicates inner mucus layer. **F)** and **H)** (thickness of inner mucus layer) Unpaired t-test; **G)** and **H)** (KLF4 staining) unpaired t-test with Welch’s correction; results are shown as mean ± SD; * p ≤ 0.05, ** p ≤ 0.01.

## DISCUSSION

Within the colonic crypt, mitochondria maintain the energy gradient, which is necessary for efficient cell differentiation and proliferation and thereby critical in determination of IEC fate (2, 10, 12). Mitochondrial disturbance and dysfunction of goblet cells are hallmarks of UC pathology (3-6, 13), which presents as a multifactorial disease, where inflammation is caused by a disruption of the colonic epithelial and mucus barrier. Terminally differentiated goblet cells have a pivotal role in the maintenance of intestinal barrier integrity and their differentiation is presumably regulated by a metabolic switch from glycolysis to mitochondrial OXPHOS (2, 17). In order to understand the molecular basis and disease origin of UC, it is necessary to find the underlying cause of mitochondrial dysfunction and to unravel a potential link to impaired goblet cell function.

Both IBD subtypes, UC and CD, are disorders of the gastrointestinal tract, which display dysfunctional mitochondria. Nevertheless, while mitochondrial disturbance results in aberrant development of Paneth cells in CD (35), we here present abrogation of goblet cell differentiation through insufficient mitochondrial respiration as a potential cause for disease development in UC. Impaired induction of goblet cell differentiation in inflamed UC but not CD has been previously reported (8). Our data indicate that defective terminal differentiation of goblet cells is already present in non-inflamed colonic tissue of UC patients in remission, defined by diminished mucus filling of goblet cells and reduced expression of terminal goblet cell differentiation marker *KLF4* compared to non-IBD controls. In line with this tenet, reduced numbers of goblet cells and a defective colonic mucus layer enabling bacterial invasion were already published for UC patients in remission (6, 7, 9, 36). While the reason for mitochondrial dysfunction in CD is still unknown, data presented here suggest that loss of p32, which is postulated to be the main driver of OXPHOS, is the underlying cause of metabolic dysfunction and secondarily of defective goblet cell function in UC. Interestingly, p32 has been previously proposed to be dysregulated before disease onset of UC but not CD, strengthening its role as a potential causative factor in disease development (37).

In addition to the observation that low p32 expression is accompanied by mitochondrial dysfunction and defective goblet cell maturation in UC, we present experimental evidence that induction of goblet cell differentiation is dependent on p32-regulated mitochondrial function *in vitro*. Stimulation of a mucus producing goblet cell-like cell line with the SCFA butyrate resulted in induction of OXPHOS and terminal differentiation. Of main interest, differentiation was abolished by *p32* silencing and mucus secretion was impaired after treatment with OXPHOS inhibitors. In line with these observations, polymorphisms in nuclear encoded mitochondrial genes involved in ATP generation, namely uncoupling protein 2 (*UCP2*) and *SLC22A5*, encoding the organic carnitine transporter 2 (OCNT2), have been described as risk factors for UC (38, 39). In addition, inhibition of intestinal fatty acid ß-oxidation as well as genetic ablation of *UCP2* or *OCNT2* in mice resulted in experimental colitis (3, 40, 41). Conversely, conplastic mice with high mucosal OXPHOS and ATP levels have been already demonstrated by our group to be protected from experimental colitis (42).

Apart from its role as a regulator of mitochondrial function, p32 has been described to interact with various proteins localized on the cell surface, the nucleus, the cytoplasm or the extracellular space (43). Binding of p32 to the globular heads of C1q reportedly inhibits classical pathway complement activation (21). Furthermore, interaction of p32 with serum proteins involved in blood clotting and fibrin polymerization as well as binding to various bacterial or viral antigens (44), might play a role in the prevention of intestinal inflammation. Whether low levels of p32 observed in UC might lead to impairment in any of the mentioned pathways, will be a topic of further investigations.

Here, we describe high level of KLF4-expressing terminally differentiated goblet cells as a healthy state and as necessary for mucus barrier integrity. In ATP8-mutant mice, carrying a mutation in complex V of the respiratory chain, we observed low colonic expression of p32 accompanied by reduced *klf4* mRNA expression, a diminished number of goblet cells and a thinned mucus layer. The transcription factor KLF4 specifically controls goblet cell fate, since in mice with intestinal deletion of *klf4* both colonocytes and enteroendocrine cells appear to undergo normal maturation. Additionally, cell proliferation and cell death rates appear unchanged in *klf4*-deficient mice, while goblet cell numbers are reduced by 90% (17). In general, goblet cells are recognized to be a major line of defense in the intestinal mucosa. The two-layered colonic mucus system separates bacteria from the host epithelium and the continuous self-renewal pushes bacteria out into the lumen, while animals with a penetrable mucus layer develop spontaneous colitis (36). Notably, high KLF4 levels suppress development and progression of intestinal neoplasia and colitis-associated colorectal cancer upon azoxymethane (AOM)/dextran sulfate sodium (DSS) treatment in mice (45).

Proliferation rather than differentiation of intestinal epithelial cells is highly important for tissue repair during active UC. Additionally, mitochondrial dysfunction in the colonic epithelium of patients with active UC has been reported to be accompanied by a reduction in fatty acid oxidation (13). We have recently published, that caspase-1 mediated cleavage of p32 results in a metabolic switch from mitochondrial oxidation to glycolysis, thereby shifting cell fate towards proliferation (14). In line, mice deficient for caspase-1 display defects in mucosal tissue repair, being detrimental under DSS-induced colitis, while derepression of the inflammasome complex results in enhanced repair and resistance to acute colitis (46). Here, we demonstrate caspase-1 to be indeed activated in inflamed colonic tissue sections of UC patients, accompanied by a reduction of antibody binding against p32 exon 6 and a decrease of differentiated goblet cells. Taken together, loss of p32 in non-inflamed colonic tissue appears highly problematic, due to a decrease in differentiated goblet cells and thereby impaired mucus barrier function. Meanwhile, during colitis, p32 cleavage might be a physiologic mechanism necessary for induction of rapid cell proliferation for tissue repair.

We here propose nutritional intervention as a potential strategy to improve colonic p32 expression. A westernized diet, rich in glucose, is a major environmental factor contributing to UC (47) and was found to continuously activate the NLRP3 inflammasome (48, 49). Having shown, that caspase-1 mediated cleavage of p32 boosts cell proliferation (14), we *vice versa* proposed an isocaloric glucose-free, high-protein diet to result in increased p32-mediated goblet cell differentiation. Indeed, mice receiving a GFHP diet exhibited induction of p32 protein expression in the colon, which was not due to elevated *p32* mRNA level compared to controls. In line with human and *in vitro* data, GFHP mice depicted high mucosal energy level, an increased number of KLF4-positive terminally differentiated goblet cells, and a thickening of the colonic mucus layer compared to controls. Considering this, dietary intervention appears as a promising tool to modulate p32 expression, mitochondrial function, and goblet cell differentiation in the intestine.

In conclusion, we identified a new pathway linking low colonic expression of OXPHOS-regulating p32 to mitochondrial dysfunction, defective goblet cell differentiation and impaired mucus barrier formation, frequently observed in UC. Furthermore, we present a diet low in glucose as an option to induce colonic expression of p32, opening new pathways in the preventive treatment and therapy of UC.

## MATERIALS AND METHODS

### Study cohort

Tissue biopsies from the terminal ileum and colon were obtained during endoscopy as part of regular patient management in the medical department 1, University Hospital Schleswig-Holstein Campus Lübeck, Germany. Blood samples were collected at the University Hospital Schleswig-Holstein Campus Lübeck, at the University Hospital Münster, North Rhine-Westphalia, Germany and at the University Hospital Rostock, Mecklenburg Western Pomerania, Germany. Characteristics of histologically confirmed UC patients and non-IBD controls at time of endoscopy or sample collection are listed in table 1 and 2, respectively. The control group included patients who presented for a regular check-up or underwent endoscopy due to non-IBD related reasons and presented without macroscopically and histological evidence of mucosal inflammation. Diagnosis of UC and classification of patients into remission and disease flare was based on clinical, endoscopic and histopathologic findings. Categorization into inflamed and non-inflamed tissue was solely based on histopathologic presentation. Groups were age and gender-matched. Non-IBD controls or UC patients with reported colon cancer were excluded from the study. All patients gave informed consent for sample donation and protocols were approved by the ethics committees of the University of Lübeck (0-073; 03-043; AZ 13/084A; AZ 05-112), the University of Münster (AZ 2016-305-b-S) and the University of Rostock (A 2017-0137).

### Animal experiments

All animal experiments were approved by the ethics committee, Schleswig-Holstein, Germany (C57BL/6FVB: V 242 – 63560/2017 (5-1/18); nutritional intervention: V 242 – 27664/2018 (64-5/17)). Mice were maintained at the University of Lübeck under specific pathogen-free conditions at a regular 12-hour light–dark cycle with free access to food (Altromin #1324, Lage, Germany, if not indicated differently) and water. Procedures involving animals and their care were conducted in accordance with national and international laws and regulations.

C57BL/6J mice were obtained from Jackson Laboratory (Bar Harbor, US) and bred in the animal facility of the University of Lübeck. The conplastic strain C57BL/6J-mt ^FVB/NJ^ which carries a mutation in the mitochondrially encoded ATP synthase membrane subunit 8 (ATP8-mutant) was generated as described previously (31) and was maintained by repeatedly backcrossing female conplastic offspring with male C57BL/6J mice. Here, basal 2.5 to 4 months old male ATP8-mutant mice and corresponding C57BL/6J controls (B6-wt) were sampled in three independent rounds. Due to differences in basal mRNA expression of targets of interest, expression data was normalized to average B6-wt target expression for each individual experiment.

Glucose free high protein (GFHP) and isocaloric control (ctrl) diet were purchased from Ssniff (Soest, Germany). Compositions of corresponding diets are specified in **supplementary table 1**. C57BL/6 mice were ordered at an age of 7-8 weeks from Charles River (Wilmington, Massachusetts, US), were left to acclimatize on a standard chow diet until an age of 20 weeks and were than randomly distributed into GFHP-diet and isocaloric ctrl diet receiving groups. Mice were kept on the corresponding diet on an average of 70 days before sampling. Food consumption and body weight were measured once a week. Dietary intervention was performed in two independent experimental rounds.

### Cell culture

The human colorectal carcinoma cell lines HT29-MTX-E12 (Sigma Aldrich, St. Louis, US) and DiFi (50) were kept in DMEM medium supplemented with or without 1% non-essential amino acids, respectively. The human colorectal carcinoma cell line T84 was kindly provided by Markus Huber-Lang, University Hospital Ulm, Baden-Wuerttemberg, Germany and grown in DMEM/F12 1:1 Medium containing 1.5% HEPES. All cell culture media were supplemented with 10% (v/v) heat-inactivated FCS, 100 U/ml penicillin, and 100 mg/ml streptomycin. Cells were incubated at 37 °C and 5% CO_2_ in a humidified incubator. Cells were cultivated up to a maximum of 20 passages and confirmed to be negative for mycoplasma contamination every three months and when freshly thawed.

For terminal differentiation, HT29-MTX cells were either grown post confluent for 9 days as described previously (30) or stimulated with 1.25 mM butyrate (Merck Millipore, Burlington, Massachusetts, US) for 72 h in the presence or absence of 1 µg/ml LPS-EB ultrapure (InvivoGen, San Diego, California, US). 2,4-Dinitrophenol (DNP; SantaCruz, Dallas, Texas, US) or Oligomycin (Agilent, Santa Clara, California, US) were applied at indicated concentrations for 24 hours to inhibit mitochondrial respiration. Further, HT29-MTX cells were transiently transfected with 50 µM silencing RNA (siRNA) specific for human *p32* (exon 3; s2138; Thermo Fisher Scientific, Waltham, Massachusetts, US) or control siRNA (Thermo Fisher Scientific) by reverse lipofection using Lipofectamine 3000 reagent (Thermo Fisher Scientific) for 96 hours or were left untreated. After 24 hours, cells were stimulated with 1.25 mM butyrate for 72 hours or were left untreated.

### RNA extraction, cDNA synthesis and quantitative real-time PCR

Isolation of total RNA from tissue biopsies or cell pellets was performed using the innuPREP RNA Mini Kit (Analytic Jena AG, Jena, Germany) according to manufacturer’s guidelines. Additional DNA digestion was performed two times after binding of RNA to RNA-column with 4 units DNase (Sigma Aldrich) in according reaction buffer for 20 min at RT. For cDNA synthesis, 1 µg of isolated RNA was transcribed with 100 pmol Oligo(dt)18 (Metabion, Steinkirchen, Germany), 20 U RiboLock RNase inhibitor (ThermoFisher Scientific), dNTP Mix (0.2 mM for each dNTP), 200 U RevertAid H Minus reverse transcriptase (ThermoFisher Scientific) in corresponding reaction buffer at 42°C for 60 min. Target amplification was performed by quantitative reverse transcriptase PCR (qRT-PCR) on the StepOne real-time system (ThermoFisher Scientific) applying Perfecta SYBR Green Supermix (ThermoFisher Scientific) and 0.5 µM forward and reverse primer. Following cycling conditions were applied: initial denaturation at 95 °C for 5 min; 40 cycles of denaturation at 95 °C for 45 sec, annealing at appropriate temperature (55 °C) for 30 sec and elongation for at 72 °C 30 sec min. Melting curve profiles were produced and data were analyzed following the 2^−dCt^ algorithm by normalized to *β-actin*. Primer sequences are listed in **supplementary table 4**.

*P32* exon expression was additionally analyzed by Taqman probes (Thermo Fisher Scientific, **supplementary table 4**) according to manufacturer’s instructions using the StepOnePlus Real-Time PCR system. The following cycling conditions were applied: preincubation at 50 °C for 2 min and 95 °C for 10 min; 40 cycles of denaturation at 95 °C for 15 sec and annealing and elongation at 60 °C for 1 min. Probe sequences are listed in supplementary table S2. Ct-Values of targets were acquired *via* the StepOne system software and normalized to *β-actin* that served as an internal housekeeping transcript *via* the 2^-dCT^ algorithm.

### SDS-PAGE and immunoblotting

SDS-PAGE and immunoblotting was performed according to standard protocols. In short, whole-protein extracts from homogenized tissue samples or cells were prepared by cell lysis in denaturing lysis buffer containing 1% SDS, 10 mM Tris (pH 7.4), phosphatase II, phosphatase III and protease inhibitor (Sigma Aldrich). Protein extracts were separated by denaturing sodium dodecyl sulfate polyacrylamide gel electrophoresis (Bio-Rad Laboratories, Hercules, California, US) under reducing conditions and transferred onto polyvinylidene difluoride membranes. After blocking, membranes were probed with specific primary antibodies followed by respective HRP-conjugated secondary antibodies. To determine similar transfer and equal loading, membranes were stripped and reprobed with an appropriate housekeeper. Proteins of interest were visualized on a ChemiDocTM XRS+ Imaging System (Bio-Rad). Applied antibodies are listed in **supplementary table 5**.

### Histology and microscopy analyses

Immunohistochemical (IHC) staining in paraformaldehyde (PFA)-fixed and paraffin-embedded tissue biopsies was performed according to standard protocols. After deparaffinization, rehydration, endogenous peroxidase blockage and antigen retrieval, tissue slides were probed with specific primary antibodies or isotype control antibodies, followed by respective HRP-conjugated secondary antibodies or HRP-labelled polymers (both listed in **supplementary table 5**). Tissue slides were incubated with DAB-substrate (Dako, Jena, Germany) and counterstained with Mayer’s hemalum solution or Alcian-blue. Images were obtained and analyzed on an Axio Scope.A1 microscope (Zeiss, Oberkochen, Germany) utilizing the ZEN imaging software (Zeiss). If appropriate, stained areas were quantified *via* the color deconvolution plugin for the software ImageJ (51).

For Muc2 immunofluorescent staining and quantification of mucus layer thickness, colonic biopsies were fixed in Carnoy’s solution before paraffin-embedding. Slides were probed with specific antibodies for murine Muc2 or according isotype control, followed by incubation with respective fluorochrome-labelled IgG secondary antibody and counterstaining using DAPI (Sigma-Aldrich). Applied antibodies are listed in **table 5**. Mucus layer thickness was measured at least at four different representative positions per slide per animal using the AxioCam software.

### ELISA

For detection of extracellular Muc5AC by ELISA pure supernatant from cells was coated at 4 °C overnight. Intracellular protein was detected in native protein isolates, coated with 50% coating buffer, containing 0.3% w/v Na_2_CO_3_ * 10 H_2_O and 0.6% w/v NaHCO_3_, pH 9.6. After blocking, Muc5AC was detected using a Muc5AC-specific primary antibody in combination with a respective HRP-conjugated secondary antibody listed in **supplementary table 5**. Optical density was measured at 450 nm against a reference wavelength of 540 nm on a SpectraMax iD3 microplate reader (Molecular Devices, San José, California, US).

### Seahorse assay

For determination of OCR and ECAR via Seahorse assay, 5×10^3^ HT29-MTX cells were seeded in a Seahorse XF24 cell culture plate in DMEM medium containing 5 mM glucose, 1% non-essential amino acids, 10% (v/v) heat-inactivated FCS, 100 U/ml penicillin, and 100 mg/ml streptomycin. Cells were stimulated with 1.25 mM butyrate for 72 hours or were left untreated. OCR and ECAR was determined in standard Seahorse medium on day three after seeding before and after injection of 2 µM oligomycin on a XF24 analyzer (Agilent) according to manufacturer’s instructions.

### Lactate assay

L-lactate levels were measured in serum or plasma samples (1:5 diluted in PBS) and in cell culture supernatants (1:10 diluted in PBS) according to manufacturer’s instructions (Megazyme, Wicklow, Ireland). Lactate level in cell culture supernatant were normalized to cell count.

### Statistics

Statistical analysis was performed using the GraphPad Prism version 6 (San Diego, California, US). Outliers were identified by Grubbs’ test (significant level α = 0.05). The F test was used to compare variances and D’Agostino–Pearson test was applied to test for normal distribution. Statistical differences between two groups were analyzed by unpaired t-test or paired t-test (normally distributed data), unpaired t-test with Welch’s correction (significant different variances) or Mann–Whitney U-test (not-normally distributed data). For comparison of more than two groups one-way analysis of variances (ANOVA) with Bonferroni post-test was applied. Uncorrected Fisher’s Least Significant Difference test was employed for data sets with two variables. Correlation analysis was performed by obtaining the Spearman’s rank correlation coefficient. P-values were calculated and null hypotheses were rejected when p ≤ 0.05. Data are shown as mean with 95% confidence interval (mean ± 95% CI), as mean ± standard deviation for small data sets (mean ± SD) or median with interquartile range for data sets with large variances.

## AUTHOR CONTRIBUTIONS

CS and SD designed the concept of the study and supervised it. HS, AB, DB and SP collected and provided human biopsy samples. MiH and SI were in charge of breeding of conplastic B6-mtFVB mice. AGS, EIP and BG provided expertise on p32 and an antibody against p32 exon 6. AS, MaH, HeS, KS, MR and AR performed the experiments and acquired the data. AS and SD analyzed and interpreted the data. AS, SD and CS drafted the article. All authors read and approved to the final manuscript.

### ACKNOWLEDGMENTS

The authors thank all the participating patients for agreeing to support this study, Prof. Jan Rupp for providing the Seahorse XF24 analyzer from Agilent and Prof. Huber-Lang for sharing the T84 cell line.

This work was supported by the German Research Foundation (Research grants SI 1518/3-1 to CS and DE 1874/1-2 to SD), the National Institutes of Allergy and Infectious Diseases, US (R01 AI 060866, R01 AI-084178, and R56-AI 1223476 to BG) and the NIH/NCI cancer support grant, US (P30 CA008748 to the Memorial Sloan-Kettering Cancer Center). CS is Fresenius Kabi endowed professor for Nutritional Medicine.

